# Task-related motor response inflates confidence

**DOI:** 10.1101/2020.03.26.010306

**Authors:** Marta Siedlecka, Borysław Paulewicz, Marcin Koculak

## Abstract

Studies on confidence in decision-making tasks have repeatedly shown correlations between confidence and the characteristics of motor responses. Here, we show the results of two experiments in which we manipulated the type of motor response that precedes confidence rating. Participants decided which box, left or right, contained more dots and then reported their confidence in this decision. In Experiment 1, prior to confidence rating, participants were required to follow a motor cue. Cued-response type was manipulated in two dimensions: task-compatibility (the relation between response set and task-relevant decision alternatives), and stimulus-congruence (spatial correspondence between response key and the location of the stimulus that should be chosen). In Experiment 2, a decision-related response set was randomly varied in each trial, being either vertical (task incompatible) or horizontal (task-compatible, spatially congruent and incongruent). The main results showed that choice confidence increased following task-compatible responses, i.e. responses related to the alternatives of the choice in which confidence was reported. Moreover, confidence was higher in these conditions, independently of response accuracy and spatial congruence with the ‘correct’ stimuli. We interpret these results as suggesting that action appropriate in the context of a given task is an indicator of successful completion of the decision-related process. Such an action, even a spurious one, inflates decisional confidence.

## INTRODUCTION

Metacognition consists of cognitive processes that monitor and control lower-order cognitive processing (Koriat & Shitzer-Reichert, 2002; Nelson & Narens, 1994). Metacognitive judgments such as decision confidence, judgments of learning, or visibility ratings are the results of these processes. It has been proposed that such judgments are based on assessment of the evidence leading to a given decision or recall (Higham, Perfect, & Bruno, 2009; Kiani & Shadlen, 2009; Vickers & Lee, 1998), but also on internal and external cues such as cognitive experiences or task-related knowledge (e.g., Bjork, Dunlosky, & Kornell, 2013; Koriat & Ma’ayan, 2005). In recent years there has been a growing amount of data suggesting that confidence in decisions is the result of metacognitive assessment of the whole process that leads to a decision and the relevant motor response (Boldt & Yeung, 2015; Filevich, Koß, & Faivre, 2020; Fleming, Maniscalco, Ko, Amendi, Ro, & Lau, 2015; Gajdos, Fleming, Saez Garcia, Weindel, & Davranche, 2019; Kiani, Corthell, & Shadlen, 2014; Siedlecka, Hobot, Skóra, Paulewicz, Timmermans, & Wierzchoń, 2019; Wokke, Achoui, & Cleeremans, 2019). In the study reported in this article, we aimed to test whether confidence judgments are affected by certain response characteristics that could be monitored in the process of action preparation and execution.

Studies on confidence in decision-making tasks have consistently shown correlations between confidence level and action characteristics. Confidence in incorrect or slow responses is typically lower than confidence in correct and fast responses (Faivre, Filevich, Solovey, Kühn, & Blanke, 2018; Gajdos et al., 2019; Kelley & Lindsay, 1993; Kiani et al., 2014; Koriat & Ma’ayan, 2005; Siedlecka, Skóra, Paulewicz, Fijałkowska, Timmermans, & Wierzchoń, 2019b; Wokke et al., 2019). It has also been shown that confidence level varies gradually with the magnitude of post-response error-related electroencephalographic activity (Boldt & Yeung, 2015; Scheffers & Coles, 2000). Another study has shown that confidence is higher when subthreshold motor preparatory activation is present before a response, even when this activation is not associated with a correct action (Gajdos et al., 2019). Fleming and colleagues (2015) showed that confidence decreases when the action-selection process is disrupted. They used transcranial magnetic stimulation (TMS) to randomly stimulate the left or right dorsal premotor (PMd) cortex of participants who performed perceptual decision tasks using their left or right hand to respond. Lower confidence was observed in trials in which participants chose a response that was incongruent with the side of premotor area stimulation.

The results of experimental studies have shown that metacognitive judgments are affected by manipulating the requirement to respond. Confidence judgments predict performance better when they are assessed after rather than before a decision-related response (Siedlecka, Paulewicz, & Wierzchoń, 2016; Siedlecka et al.,, 2019b; Wierzchoń, Paulewicz, Asanowicz, Timmermans, & Cleeremans, 2014; Wokke et al., 2019, although see: Charles, Chardin, & Haggard, 2020; Filevich, Koß, & Faivre, 2020) and they are more accurate when they follow one’s own action compared to observed actions of others (Pereira, Faivre, Iturrate, Wirthlin, Serafini, Martin, Desvachez, Blanke, Van De Ville, & Millán, 2018). In our previous study, action involving the same response set as a decision-related response increased reported metacognitive awareness (Siedlecka et al., 2019a). In the aforementioned study, participants were asked to perform a perceptual discrimination task, and the stimulus was immediately followed by a cue requiring an action that was irrelevant to the main task but could be the same, opposite, or neutral to the correct stimulus-related response. Specifically, participants were asked to determine the orientation of Gabor gratings (left or right), and then a cue would require either a “left”, “right” or “neutral” (pressing the space key) response. The results showed that participants reported a higher level of stimulus awareness after carrying out a response that was either congruent or incongruent with the correct response, compared to the neutral one. We interpret these findings as supporting the hypothesis that a spurious motor response that overlaps potential responses to the stimuli is an indicator of successful decision process completion, and therefore it inflates subjective certainty.

The aim of this study is to further investigate the role of action in confidence judgment by testing how confidence level is affected by different types of preceding motor responses. On one hand, the previous evidence suggests that confidence could be increased by the mere presence of a decision-related action. On the other hand, metacognitive judgments might reflect the results of performance monitoring, a process that is thought to detect errors and response conflicts and thus enable action control and regulation (Botvinick, Cohen, & Carter, 2004; Ridderinkhof, Ullsperger, Crone, & Nieuwenhuis, 2004). Response conflicts are studied in action control research and are assumed to stem from interference between different motor tendencies. They may occur at different levels of information processing, such as stimulus encoding, response selection, or response execution (van Veen, Cohen, Botvinick, Stenger, & Carter, 2001). The incongruence appears when two different aspects of the same stimuli are linked to different motor responses, or when a stimulus linked to a specific response is followed by a prime linked to a competitive motor plan (Desender, Van Opstal, & Van den Bussche, 2017). In such cases, a congruency effect is observed: responses are slower and more often erroneous in incongruent trials compared to congruent ones. Detection of conflict is thought to trigger regulatory processes (e.g. Botvinick et al., 2004; Gratton, Coles, & Donchin, 1992). Some studies suggest that cognitive control can be achieved automatically and unconsciously (Charles, Gaillard, Amado, Krebs, Bendjemaa, & Dehaene, 2017; Lau & Passingham, 2007; Van Gaal, Ridderinkhof, Fahrenfort, Scholte, & Lamme, 2008), and the results of conflict monitoring do not have to necessarily translate into metacognitive experience. No direct relations between confidence and cognitive conflict have been detected so far; however, it has been shown that conflicts might lead to a subjective feeling of difficulty (Desender et al., 2017; Morsella, Gray, Krieger, & Bargh, 2009; Morsella, Wilson, Berger, Honhongva, Gazzaley, & Bargh, 2009). Some authors even suggest that experience of conflict is necessary for subsequent regulation (Desender, Van Opstal, & Van den Bussche, 2014; Questienne, Van Opstal, van Dijck, & Gevers, 2016).

In the presented study, we have introduced different types of response conflict. We used a perceptual discrimination task in which participants were required to decide which box (left or right) contained more dots and to report their decision confidence. In Experiment 1, confidence judgment was preceded by a motor response to a cue presented on the screen. We manipulated the spatial congruence between the cued response and the position of the stimuli that should be chosen; we also manipulated the compatibility between a cued response and task-relevant decision alternatives. In Experiment 2, we manipulated the spatial congruence between the response and “correct” stimuli by manipulating the response set. The main results showed that choice confidence is increased following an action that is compatible with the decision-relevant dimension (left/right) compared to the control conditions.

## EXPERIMENT 1

Our previous study (Siedlecka et al., 2019a) showed that directional motor responses, that is responses compatible with task-relevant stimuli characteristics (left/right), increase metacognitive ratings of stimuli awareness compared to the neutral condition, even if they are spatially incongruent with the stimulus that should be chosen. However, participants in this task responded to the stimulus using only one hand, which could have reduced the experienced conflict between the position of the stimuli (and therefore prepared action) and action forced by the cue. In Experiment 1, we aimed to replicate our previous results. We introduced some methodological changes, including better manipulation and control of spatial response conflict, and we collected confidence ratings instead of perceptual awareness judgments.

In Experiment 1, participants were asked to decide which of the two boxes presented on the left or right of the screen contained more dots; they were then asked to report their choice and confidence. However, immediately after stimulus presentation and before task-related responses were provided, participants were required to respond to the motor cue that was irrelevant to the main task. We used four motor cues: left arrow, right arrow, arrow pointing left and down, arrow pointing right and down. In this way, we created two types of compatibility: general spatial congruence between the cued response and the task-relevant dimension of the stimuli, and task compatibility between the cued response and the allowed response to the stimuli. This resulted in four conditions: task-compatible and stimulus-congruent (TC-SC), task-compatible and stimulus-incongruent (TC-SI), task-incompatible and stimulus-congruent (TI-SC), and task-incompatible and stimulus-incongruent (TI-SI). For example, a TC-SI trial means a participant is cued to press the right arrow key when there are more dots on the left side; a TI-SC trial is when a person is required to press the left-down key when there are more dots on the left side. We assumed that SI trials would create conflict between motor preparation in response to the stimulus and response to the cue, while TI trials create conflict between allowed responses to the stimulus and cued response. We hypothesized that decision accuracy would be affected mainly by spatial congruence, but confidence would be affected by both compatibility and congruence in an additive way: it would be lowest in task-incompatible and stimulus-incongruent trials (TI-SI) and highest in task-compatible and stimulus-congruent trials (TS-SC).

### Methods

#### Participants

Forty-eight participants (7 males) aged 19–36 (M = 21.04, SD = 3.57) took part in the experiment in return for course credits. All participants had normal or corrected-to-normal vision and gave written consent to participation in the study. The ethical committee of the Institute of Psychology of Jagiellonian University approved the experimental protocol.

#### Materials

The experiment was run on computers using PsychoPy software (Pierce, Gray, Simpson, MacAskill, Höchenberger, Sogo, Kastman, & Lindeløv, 2019) with LCD monitors (1920 × 1080 resolution, 60 Hz refresh rate). Stimuli consisted of two frames containing a number of dots (Bold & Yeung, 2015). The frames were placed on the right and left side of the fixation point, and they were 160 × 160 px large, resulting in a visual angle of 3.8° when viewed from 60 cm. Each frame contained dots arrayed in a 10-by-10 matrix. The displays were randomly generated for each trial and each subject. The sum of the dots between the two fields was always 100. During the first training session, one field contained 30 or 40 dots, while the other contained 70 or 60 dots. During the second training session, another level of difficulty was added: 45 dots on one side and 55 dots on the other side. During the experimental session, each frame contained 47 or 53 dots. The difficulty level for the experimental session was set through the piloting of the task, aiming for about 80% accuracy.

To measure participants’ decision accuracy and their confidence, we used an eight-point scale: “Left – I am very confident”, “Left – I am quite confident”, “Left – am not confident”, “Left – I am guessing”, “Right – I am guessing”, “Right – I am not confident”, “Right – I am quite confident” and “Right – I am very confident”. Participants used the keys on the left and right side of the keyboard (q, w, e, r, u, i, o, p, respectively), which were labelled with colors and numbers: “I am guessing” answers were covered with white stickers with the letter L or R; “I am not confident” with yellow stickers with number 2; “I am quite confident” with orange stickers with number 3; and “I am very confident” with red stickers with number 4 (Figure 1).

**Figure 1.**
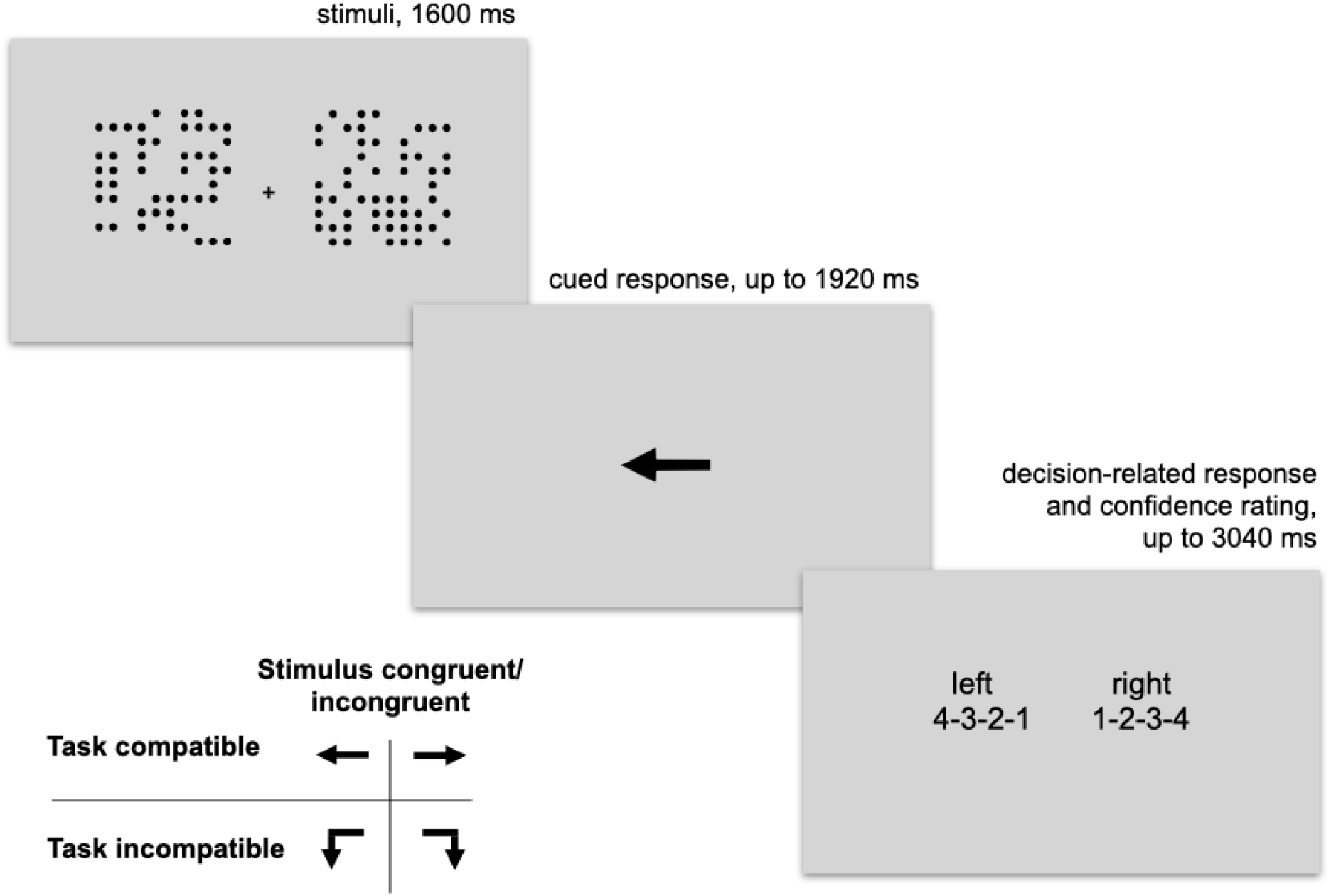
The outline of the procedure in Experiment 1.

#### Procedure

The experiment consisted of a series of trials in which participants performed a perceptual decision task and simultaneously rated their subjective confidence in this decision. The perceptual task required participants to judge which of two briefly flashed frames contained more dots. Additionally, before responding to the perceptual decision, task participants were required to respond to the motor cue presented on the screen. Participants were told that this task was irrelevant to the main task, but they should respond to the cues as quickly and accurately as possible. There were four types of cues: arrow left, arrow right, arrow down and left, arrow down and right. Arrows pointing left and right required the “L” and “P” responses, respectively (task-compatible condition), while the down arrows required responses with the keys labelled X and Y, respectively (task-incompatible condition).

Participants were tested in a computer laboratory in small groups. They were told they would take part in an experiment about decision-making. They were asked to place their left hand on the left side on the keyboard, so their four fingers stayed on the keys labelled 4-3-2-L and their thumb stayed on the key labelled X. Similarly, their right hands were on the right side of the keyboard, the keys labelled R-2-3-4, and their thumb was on the key labelled Y.

The experiment started with three training sessions. During the first session (10 trials), participants were presented with stimuli for 320 ms and were only asked to decide which frame contained more dots (using the L and R key). In the second session (12 trials), the arrows and scale were added; the third session (8 trials) was exactly like the following experimental session.

During the experimental block, each trial started with a fixation point for 800 ms, followed by the presentation of the dots that were on the screen for 160 ms. Immediately afterwards, the motor cue was presented for 1920 ms. Participants were asked to respond within this time; they used their left index finger to respond to the arrow pointing left; left thumb to respond to the arrow pointing left and down; right index finger to respond to the arrow pointing right; right thumb to respond to the arrow pointing right and down. Immediately after the motor cue disappeared or participants responded, the scale was presented on the screen for 3040 ms. Participants were asked to decide if there were more dots on the left or right side and simultaneously rate their confidence. Participants used their left hand if they decided there were more dots on the left (keys 4-3-2-L), and their right hand if they decided there were more dots on the right (keys R-2-3-4). The outline of the procedure is presented in Figure 1.

There were 480 experimental trials separated by short breaks, and one longer break in the middle of the experiment during which participants performed light physical activity and were offered candies.

#### Sample size estimation

In our previous study, the effect of compatibility on confidence was estimated at around 0.15 (Siedlecka et al., 2019). Because of the procedural differences between the experiments, we used the lower 95% confidence interval (0.06) as a conservative estimate of the expected effect size. Since the number of trials per condition was similar between the experiments, we assumed that the magnitude of standard errors should also be similar. In order to reliably detect an effect size of 0.06 we need standard errors of 0.03, which in this case is 3/4 of the size of those obtained in the previous study (0.04), meaning that the sample size should be around 40 (4/3 squared times previous sample size). However, we typically overbook the number of participants in case some of them do not turn up, perform poorly, or technical problems occur.

### Results

Statistical analyses were run in the R statistical environment using linear or logistic mixed regression models. The models were fitted using the lme4 and lmerTest packages (Bates, Maechler, Bolker, & Walker, 2015; Kuznetsova, Brockhoff, & Christensen, 2017; R Core Team, 2019).

#### Data selection and descriptive statistics

Prior to data analysis, we removed all non-complete trials, i.e. those in which at least one response was missing (confidence rating or cued response, in total 1% of all trials). Also, some participants were omitted from further analyses: one that used only one scale point; one that had a decision accuracy of 53%; and four that had a cued-response accuracy below 60%. The cutoff point was based on a Bayesian mixture model fitted to participants’ accuracy (see e.g. Lee & Wagenmakers, 2014); this was based on the assumption that each participant came from one of two distributions that differed in the probability of any given trial being correct.

The data of forty-two participants were included in the main analyses. The average decision accuracy was 76%. Accuracy in each condition is presented in Table 1. Since we were interested only in trials in which participants followed the motor cue, all trials with incorrect responses to the cue were removed prior to further analysis (515 trials).

**Table 1.** Decision accuracy, cued-response accuracy, reaction time (ms) for correct cued responses, reaction time for decision and confidence responses, and average confidence for all four conditions: TC-SC (task compatible and stimulus congruent), TC-SI (task compatible and stimulus incongruent), TI-SC (task incompatible and stimulus congruent, and TI-SI (task incompatible and stimulus incongruent).

| Condition | Decision-related response accuracy | Cued-response accuracy | Cued-response RT (SD) | Confidence & decision RT (SD) | Average confidence (SD) |
| --- | --- | --- | --- | --- | --- |
| TC-SC | 81% | 97% | 783 (322) | 474 (340) | 2.69 (0.93) |
| TC-SI | 70% | 98% | 826 (333) | 474 (345) | 2.66 (0.93) |
| TI-SC | 80% | 98% | 800 (295) | 457 (315) | 2.62 (0.94) |
| TI-SI | 73% | 97% | 824 (300) | 451 (327) | 2.60 (0.92) |

#### Confirmatory analyses

##### The effect of response type on decision accuracy

Confidence level is usually strongly related to decision accuracy, therefore we tested how accuracy differed between conditions. The logistic mixed model (with random intercept) showed that decision accuracy is affected by congruence between stimulus and cued response: accuracy is higher in SC compared to SI trials (*z =* −14.66, *p* < .001). We did not detect such an effect for task compatibility (*z =* 1.07, *p* = .28). However, another model in which an interactive term is included showed a strong interaction between two types of compatibility: in the task-incompatible condition, the effect of stimulus-congruence on accuracy is lower compared to the task-compatible condition (*z* = 2.9, *p* <.001).

##### The effect of response type on confidence level

We tested how confidence level is influenced by the two types of variables: task-compatibility and stimulus-congruence. First we used a linear mixed model with random intercept to find out whether the two variables interact; it showed that the interactive effect of both types of compatibility was not significant. The model with this interactive term was not significantly better than the model without it (chisq(1) = 1.9, *p* = .17), so we fitted an additive model with a random intercept (Table 2). The model fit was significantly better than that of the null model (chisq(2) = 35.6, *p* < .001). The results show that both types of compatibility affect confidence level in an additive way. Participants were more confident in their decisions in TC trials compared to TI trials, and they were more confident in SC trials compared to SI trials.

**Table 2.**
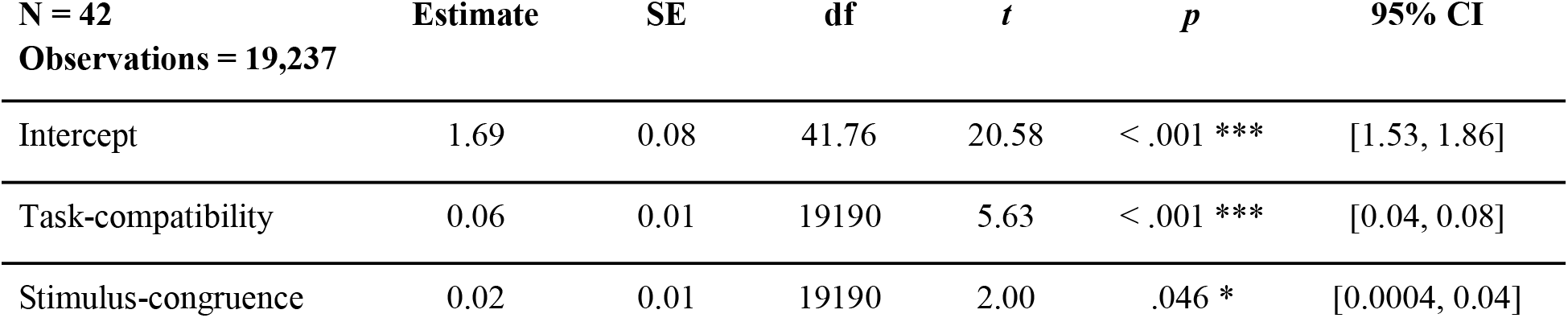
The linear mixed model estimating the effect of task-compatibility and stimulus-congruence on confidence ratings.

| N = 42<br>Observations = 19,237 | Estimate | SE | df | *t* | *p* | 95% CI |
| --- | --- | --- | --- | --- | --- | --- |
| Intercept | 1.69 | 0.08 | 41.76 | 20.58 | < .001 *** | [1.53, 1.86] |
| Task-compatibility | 0.06 | 0.01 | 19190 | 5.63 | < .001 *** | [0.04, 0.08] |
| Stimulus-congruence | 0.02 | 0.01 | 19190 | 2.00 | .046 * | [0.0004, 0.04] |

However, since one can expect that the difference in confidence between the stimulus-congruent and the stimulus-incongruent conditions could at least partially stem from differences in decision accuracy, we added accuracy (as a covariant) and random effect of accuracy to our model. The results show that although the effect of task compatibility on confidence remains, the effect of stimulus congruence is no longer significant (Table 3).

**Table 3.**
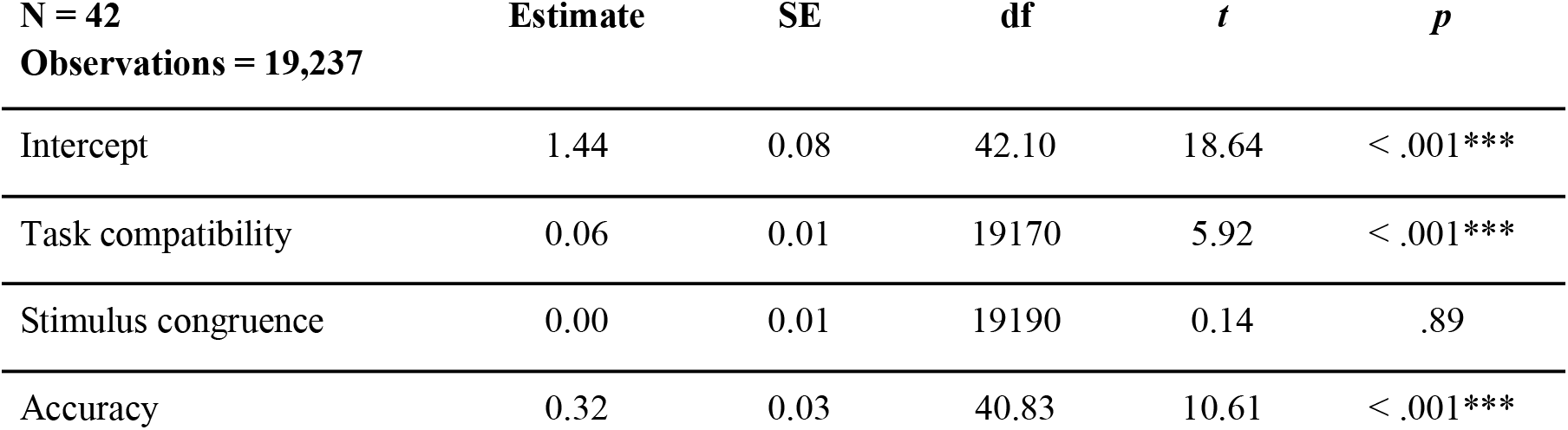
The linear mixed model estimating the effect on confidence ratings of task compatibility, stimulus congruence, and response accuracy.

| N = 42<br>Observations = 19,237 | Estimate | SE | df | *t* | *p* |
| --- | --- | --- | --- | --- | --- |
| Intercept | 1.44 | 0.08 | 42.10 | 18.64 | < .001*** |
| Task compatibility | 0.06 | 0.01 | 19170 | 5.92 | < .001*** |
| Stimulus congruence | 0.00 | 0.01 | 19190 | 0.14 | .89 |
| Accuracy | 0.32 | 0.03 | 40.83 | 10.61 | < .001*** |

The next analysis aimed to test whether the difference in confidence level between TC and TI conditions depends on decision accuracy or whether participants are more confident in the task-compatible condition for both correct and incorrect responses. A linear mixed model with random effects of accuracy, compatibility, and their interaction revealed that the difference between correct and incorrect decisions in the TC condition did not change significantly in the TI condition (*t*(>100) = 0.03, *p* = .98). Therefore, we can say that task compatibility increases confidence in both correct and incorrect decisions.

#### Exploratory analyses

##### How confidence is related to congruence between cued-response and chosen response

Finally, we tested whether response accuracy interacts with the SC condition. Please note that – independently from manipulating stimulus-congruence – we can measure response congruence, i.e. the relation between a cued response and an actual decision-related response (here, indicating whether there are more dots on the left or right side). Incorrect responses in the SC condition are actually congruent with cued-motor responses, and incorrect responses in the stimulus-congruent condition are actually congruent with cued-motor responses. If confidence is related to congruence between cued responses and the chosen response to a stimulus, we could expect higher confidence in correct SC and incorrect SI trials compared to incorrect SC and correct SI trials. However, a model with an interactive term (and random effect of intercept, congruence and accuracy) did not reveal a statistically significant interaction: t(>100) = 0.9, *p* = .37.

#### The effect of cued-response type on response times

To find out how compatibility and congruence affected the time of cued responses and confidence ratings, we used linear mixed models with a random intercept effect. The contrast analysis did not reveal significant differences between the four conditions in cued-response latencies (*p* > .18) or decision and confidence RTs (*p* > .5).

### Discussion

The results of this experiment confirmed the hypothesis that confidence level is affected by the preceding action. We manipulated the type of the motor response in two dimensions: stimulus congruence, i.e. the spatial relation to the location of the to-be-chosen stimuli; task-compatibility, i.e. the overlap with the response set that is associated with task-related decision alternatives. The results show that – independently of stimulus-congruence – confidence is higher following task-compatible motor responses compared to task-incompatible responses. Stimulus-congruence affects decision accuracy, but we did not detect its influence on confidence judgment when response accuracy was controlled for. Moreover, task-compatible trials increase confidence in both correct and incorrect decisions. Altogether, we interpret the results as suggesting that decision confidence is inflated after spurious motor responses, i.e. responses that are compatible with task-related responses, even if this response spatially conflicts with the correct response to the stimulus.

These results replicate the previous finding which showed that metacognitive awareness increases after task-compatible cued responses (Siedlecka et al., 2019a). The results are consistent despite a number of methodological changes, such as the type of metacognitive judgment, the measurement scale, and the way participants responded (i.e. using one or two hands).

However, one alternative explanation for the results is that it was not the response itself but the motor cue presentation (and its congruence with correct stimulus) that influenced confidence judgments. In the next experiment, we aimed to remove this possible confounder. Additionally, we simplified the task such that it did not require participants to carry out two discrimination tasks in addition to the confidence rating, and they had to store stimuli-related decision in memory during cued-response tasks.

## EXPERIMENT 2

In Experiment 2, participants were required to provide their decision-related response immediately after stimulus presentation, but this time the response key was randomly manipulated in each trial. Half of the participants were asked to decide whether there were more dots on the left side, and the other half decided whether there were more dots on the right side. After the stimulus had disappeared, an available response set was presented: either two horizontal arrows (left and right) or two vertical ones (top and down). One of the arrows was associated with a “yes” response and the other with a “no” response. Therefore, we created conditions in which the response set is compatible with the task-relevant dimension of the stimuli (horizontal response set) and is incompatible with this dimension (vertical response set). Task-compatible responses could be congruent or incongruent with the location of the correct stimuli, while task-incompatible responses were always neutral in this respect. The incongruence occurs in trials in which correct response to the stimuli is incongruent with the location of the to-be-chosen stimuli. Similarly to the results of Experiment 1, we expected response accuracy to be lower in incongruent trials than in congruent and neutral ones. We also expected that task compatibility would affect confidence, therefore confidence would be higher in both congruent and incongruent task-compatible trials compared to the neutral condition.

### Methods

#### Participants

Fifty-two participants (7 males) aged 19–46 (M = 22.67, SD = 4.44) took part in the experiment in return for a small payment (5 EUR). All participants had normal or corrected-to-normal vision and gave written consent to participation in the study. The ethical committee of the Institute of Psychology of Jagiellonian University approved the experimental protocol.

#### Materials

The dot stimulus was the same as in Experiment 1. Immediately after the stimuli presentation, the response set for each trial was presented. The response set provided information on which of the four keyboard buttons (right, left, down or up arrow) would correspond to the “yes” answer and which to the “no” answer. To simplify the task, we developed two alternative versions of the procedure: in one, participants were always asked whether there were more dots on the left side; in the other, participants were always asked whether there were more dots on the right side. Each response set cue consisted of a central letter (“L” or “R”) to remind participants about the decision they were supposed to make, and two arrows (left and right or up and down) associated with the responses “yes” and “no”. These responses were assigned randomly, i.e. each of them could be assigned to any arrow.

To measure participants’ confidence, we used a four-point scale: 1.“I am guessing”; 2. “I am not confident”; 3. “I am quite confident”; 4. “I am very confident”. Additionally, we made it possible for participants to report errors (i.e. mistakes, slips) in order to exclude cases in which they responded differently to what they intended and to better assess decision accuracy.

#### Procedure

Participants were tested in a computer laboratory in small groups. They were told they would take part in an experiment about decision-making. The experiment started with three training sessions. During the first session (10 trials), the stimuli were presented for 320 ms and participants decided which frame contained more dots (using the left and right arrows with their right hands). In the second session (10 trials), the confidence scale was added, and the third session (12 trials) was the same as the following experimental session.

During the experimental session, each trial started with a fixation point for 800 ms, followed by the presentation of the dots, which were on the screen for 160 ms. Immediately afterwards, the response set board was presented for up to 3040 milliseconds, during which participants were supposed to respond with their right hand using the arrows keys. Subsequently, the confidence scale was presented, and again participants had a maximum of 3040 milliseconds to assess their confidence in their response. Participants used the keys 1–4 and the space bar (for error reporting) on the keyboard with their left hand. There were 456 experimental trials, separated by short breaks every 30 trials, and one longer break in the middle of the experiment, during which participants performed light physical activity and were offered refreshments.

There were three experimental conditions: task incompatible (neutral in respect to the stimulus, later referred to as Neutral), task compatible and stimulus congruent (Congruent), and task compatible and stimulus incongruent (Incongruent). The vertical response set (“up” and “down”) is spatially neutral in respect to the task-related dimension, while the horizontal response set (“left” and “right”) is task compatible. Stimulus congruence was manipulated within the horizontal response set trials. The incongruent trials included, for example, cases in which there were more dots on the right side, and participants were required to press the left arrow to respond correctly (”no” when asked whether there were more dots on the left; “yes” when asked whether there were more dots on the right). The instances of congruent trials were, for example, when there were more dots on the right, and the correct response was provided by pressing the left arrow (“no” when asked whether there were more dots on the left; “yes” when asked whether there were more on the right). The outline of the procedure is presented in Figure 2.

**Figure 2.**
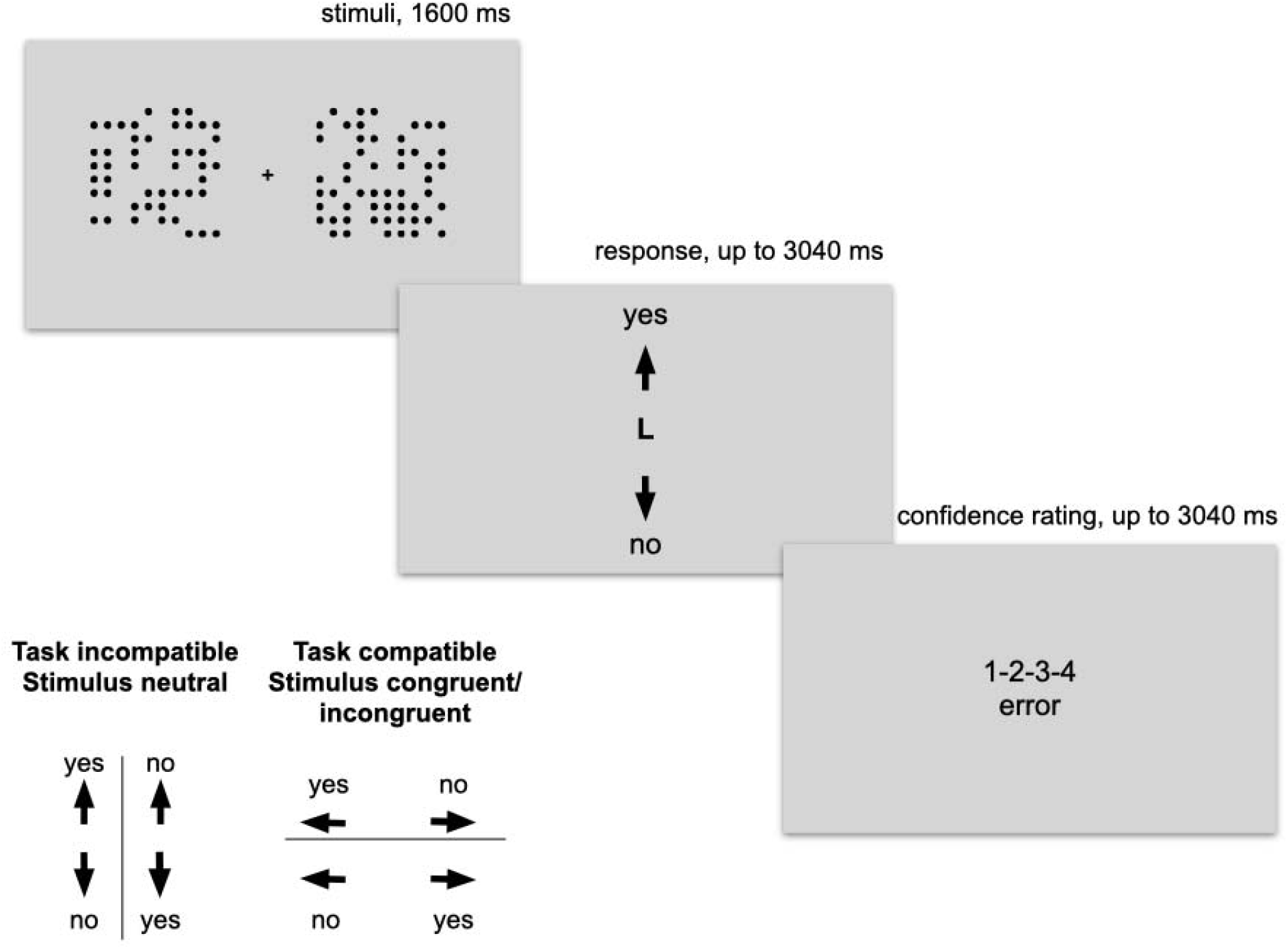
The outline of the procedure in Experiment 2.

### Results

#### Data selection and descriptive statistics

Prior to data analysis, we removed all non-complete trials, i.e. trials in which at least one response was missing (confidence rating or stimulus-related response), in total this was 1% of all trials. One participant with low scale-usage variance and seven participants with response accuracy below 55% were omitted from further analyses. We estimated standard deviation of confidence ratings for each participant; the plot of the sorted standard deviations indicated a clear discontinuity caused by one participant having a very low standard deviation (0.3).

The data of forty-four participants were included in the main analyses. The average response accuracy was 73%. We did not find significant differences between the three conditions (Neutral, Congruent, Incongruent) in accuracy (|*z*| ≤ 0.85, *p* ≥ 0.2). Similarly, we did not detect differences in d’ (|*z*| ≤ 1.05, *p* ≥ 0.3) and bias (|*z*| ≤ 0.13, *p* ≥ 0.4). Participants reported errors 349 times: in 1% of trials in the Congruent condition, and in 2% of trials in the Incongruent and Neutral conditions. We did not detect differences in the accuracy of the error reports, probably due to the very small sample size. Accuracy and other descriptive statistics are presented in Table 5.

**Table 5.** Decision accuracy, reaction time (ms) for decision and confidence responses, and average confidence for the three conditions (Congruent, Incongruent, Neutral).

| Condition | Response accuracy | Response RT<br>(SD) | Confidence RT<br>(SD) | Average confidence<br>(SD) |
| --- | --- | --- | --- | --- |
| Neutral | 73% | 1,236 (506) | 533 (385) | 2.80 (0.96) |
| Congruent | 73% | 1,160 (510) | 557 (392) | 2.83 (0.96) |
| Incongruent | 72% | 1,167 (503) | 549 (388) | 2.83 (0.97) |

#### Confirmatory analyses

##### The effect of congruency type on confidence level

In these analyses, we compared confidence level between the three main conditions: Neutral, Congruent and Incongruent. Error reports were omitted, and we included only correct responses because accuracy is related to stimulus-response congruence (i.e. the congruent condition is congruent when a given response is correct, but it becomes incongruent when a person responds incorrectly).

Linear mixed regression with random intercept and participant effect showed that confidence in the Neutral condition is lower than in the Congruent (*t* = 0.27, *p* = .02) and Incongruent conditions (*t* = 2.61, *p* = .009, Table 6). The We found no significant difference between the Congruent and Incongruent conditions (*t* = 0.28, *p* = .77).

**Table 6.** The effect of response type on confidence level.

| N = 44<br>Observations = 14,218 | Estimate | SE | df | *t* | *p* | 95% CI |
| --- | --- | --- | --- | --- | --- | --- |
| Intercept (Neutral) | 2.80 | 0.08 | 43.60 | 34.01 | <.001*** | [2.63, 2.69] |
| Congruent | 0.04 | 0.02 | 14170 | 2.27 | .02* | [0.01, 0.07] |
| Incongruent | 0.04 | 0.02 | 14170 | 2.61 | .009** | [0.01, 0.08] |

Additional models including fixed effect of accuracy and condition and random effect of accuracy and participant revealed that the effect of condition on confidence was not significantly different for incorrect responses (interactions with accuracy Neutral vs. Congruent: *t*(>100) = 0.10, *p* = .27; Neutral vs. Incongruent: *t*(>100) = 0.65, *p* = .52; Congruent vs. Incongruent: *t*(>100) = 0.40, *p* = .69).

#### Exploratory analyses

##### The relationship between “yes” and “no” responses and confidence

For exploratory reasons, we ran an analysis in order to find out whether the affirmative and negative responses were differently related to confidence level. Using the linear mixed model with random intercept and random participant effect, we did not detect a relation between confidence and the type of response (*t*(49.52) = 1.46, *p* = .15), nor did we find an interaction between response type and stimulus congruence (*t*(>100) = 1.58, *p* = .11).

##### The relationship between congruence and response times

We also compared response time between conditions: it was longest in the Neutral condition (Neutral vs. Congruent: *t*(>100) = 8.92, *p* < .001; Neutral vs. Incongruent: *t*(>100) = 7.39, *p* < .001). We found no differences between Congruent and Incongruent trials (*t*(>100) = 1.29, *p* = 0.2). Similarly, confidence rating time was longest in the Neutral condition (Neutral vs. Congruent: *t*(>100) = −3.71, *p* < .001; Neutral vs. Incongruent *t*(>100) = −3.24, *p* = .001). On the Confidence scale, no difference between response time was detected when congruent and incongruent conditions were compared (*t*(>100) = 0.39, *p* = 0.7).

### Discussion

In this experiment, we tested the effect of task compatibility and stimulus congruence on decision confidence. The results showed that participants were more confident in responses that were compatible with task-relevant decision alternatives (Congruent and Incongruent) compared to the task-incompatible condition (Neutral). We did not detect an effect of stimulus congruence or the interaction between task compatibility and response accuracy; neither did we find an effect of condition on decision task performance. Altogether, the results suggest that, compared to neutral responses, responses that correspond to task-relevant decision alternatives increase confidence.

The results support the findings from Experiment 1. In both experiments, where the task was to decide which of two boxes, left or right, contained more dots, responses involving pressing right- or left-pointing arrows were followed by higher confidence ratings, despite procedural differences and different control conditions (down-left and down-right arrows in Experiment 1 and up- and down-pointing arrows in Experiment 2).

## GENERAL DISCUSSION

The research question underlying this study concerns the relation between confidence and action. Specifically, we tested how different types of motor response affect decision confidence. In Experiment 1, we manipulated the type of irrelevant, cued motor response that followed stimulus presentation but preceded confidence rating. The cued response could be task compatible, i.e. related to the task-relevant decision alternatives (left/right), or task incompatible. At the same time, the cued response could be stimulus congruent, i.e. spatially corresponding to the location of the stimulus that should be chosen, or stimulus incongruent. Although this cued response was not directly related to decisions, task compatibility increased confidence in both correct and incorrect decisions. In Experiment 2, we varied the response key after each stimulus presentation, but this time the response directly expressed the stimulus-related decision. We manipulated task compatibility by introducing two types of response sets: vertical (task incompatible, stimulus neutral) and horizontal (task compatible, stimulus congruent and incongruent). Decision confidence was higher after task-compatible responses, irrespective of whether they were spatially congruent or incongruent with the “correct” stimuli, and whether they were correct or incorrect.

The results of both experiments are consistent, suggesting that task-related action affects the following confidence judgment. In both experiments, participants’ decisional confidence was higher after responses that related to the alternatives from which participants were required to choose. We expected these results because our previous study showed that metacognitive awareness ratings were higher after task-compatible cued responses (Siedlecka et al., 2019a). One explanation for these results is that action appropriate in the context of a given task is an indicator of successful completion of the decision-related process, and it therefore inflates confidence in one’s correct and erroneous decisions. This is consistent with the hierarchical model of self-evaluation proposed by Fleming and Daw (2017), according to which responses provide information about the decision process that is otherwise not accessible to the subject. Previous studies supporting this thesis have shown that one’s own motor response that precedes confidence judgment increases correlations between confidence and response accuracy (Pereira et al., 2018; Siedlecka et al., 2016; Siedlecka et al., 2019b; Wierzchoń et al., 2014; Wokke et al., 2019). However, in a recent study, Filevich and colleagues (Filevich et al., 2020) found that when participants were asked to continuously track decision-related characteristics of stimuli (the motion direction of the stimulus) by pressing the left and right keys, their confidence in a temporal-summation decision task was higher than in the no-tracking condition. They interpreted the results as showing that action related to a stimulus that one has to decide about provides additional sources of information, thus leading to more liberal confidence criteria.

A second way of interpreting the results is that task incompatibility reduced decision confidence. We do not find much support for this view in our data: we did not detect consistent differences in response times between the conditions, nor did we consistently observe other conflict-detection correlates such as longer response times or higher error rate (e.g. Egner, 2017). On the other hand, motor conflict occurs in situations in which one of the response tendencies is prepotent due to strong habitual associations with given stimulus-related characteristics. In our study, response-related aspects of stimuli (the side of the box which contained more dots) were probably not automatically encoded. Moreover, characteristics of the stimuli that participants decided about most likely caused preparatory activation of responses associated with both “left” and “right” decisions. Therefore, the main response conflict might have occurred in the task-incompatible condition: between two preactivated responses and the required response that is not related to a decision-relevant stimulus characteristic (i.e. “up” or “down”). This interpretation, although not supported by empirical evidence and theory, is not contradictory to the previous one. In fact, we might have observed both: increased confidence due to completion of the decision-response process, and decreased confidence due to the response conflict.

We assume that subjective reports of confidence are a result of metacognitive processing that consists of many monitoring and regulating loops that occur on consecutive stages of stimuli processing, decision making, and action preparation and execution (Paulewicz, Siedlecka, & Koculak, 2020). When looking for causal relations between target variables and metacognitive judgments, we should take into consideration that there are experimentally uncontrollable feedback loops between monitoring and regulation (Paulewicz et al., 2020). For example, in Experiment 1 the presence of the cue that created a response conflict could have increased cognitive control, which could have affected the way in which decision or action preparation was processed and monitored. In Experiment 2, the manipulation of response type affected the last stage of the decisional process, therefore it could only have influenced the subsequent confidence judgment. However, even though we manipulated the response key in Experiment 2, the response actually given by participants was determined by both their decision and the available response key, therefore we cannot rule out that confidence rating is influenced by other variables that precede motor response, even if they are hard to identify.

To conclude, although many studies have shown that confidence levels correlate with motor response characteristics (Fleming et al., 2015; Gajdos et al., 2019; Kiani et al., 2014; Boldt & Yeung, 2015; Scheffers & Coles, 2000), observational findings do not determine the causal relations between confidence and other variables. These correlations might reflect common causes of confidence, accuracy and response time, or the effect of confidence itself (i.e. the higher the confidence, the quicker the responses). Here, we show that confidence is affected by experimental manipulation of the type of motor response that precedes confidence rating. Although the idea that decision-related responses inform confidence is coherent with post-decisional models of confidence (i.e. Kvam, Pleskac, Yu, & Busemeyer, 2015; Pleskac & Busemeyer, 2010; Moran, Teodorescu, & Usher, 2015), none of those models explicitly predicts the effect of the type of action. Moreover, most confidence theories and empirical studies seem not to differentiate between decision and action. Data supporting the thesis that confidence is informed by action-related information are derived from experiments that use the same measures for decision and response. However, reaction time might reflect either the ease of the decision process or action fluency, and accuracy of response might not reflect decision accuracy if participants made a different response to the one they intended. We believe that separating at least these two stages of the decisional task by inserting spurious responses, manipulating response sets, or allowing participants to report motor errors, could improve our understanding of the influence of actions on confidence.

MS, MK, BP conceived the plan of the study. MK prepared the experimental procedure, MS ran the tests, and BP analysed the data in collaboration with MS. MS drafted the manuscript, and all the authors provided comments.

The authors would like to thank Patryk Guzda, Ewa Ilczuk and Alicja Krzyżowska for help with data collection.

